# Multiscale analysis of a non-periodic epidemic time series using wavelet transform

**DOI:** 10.1101/535146

**Authors:** Jean Gaudart, Stanislas Rebaudet, Gaetan Texier, Robert Barrais, Renaud Piarroux, Roch Giorgi

## Abstract

The aim of the present study was to develop a method for multiscale analysis of non-stationary and non-periodic epidemic time series. Indeed, the epidemiologists may need to know the features, at different resolutions, of short duration outbreaks that did not exhibit periodic cycles. Among of the large number of wavelets, we have developed a continuous wavelet that shows an analogous shape to the Haar wavelet, and leads to precise time localization. We applied the wavelet transform to the cholera epidemic, which began in October 2010 in Haiti. We determined the wavelet spectra of both the cholera case toll and rainfall time series, from September 01, 2010, to November 20, 2012 (812 days). The relationship between case toll and rainfall was analyzed using cross-wavelet spectra at different lags. Cholera case toll scalogram highlighted four epidemic bursts. Cross-wavelet analysis highlighted an absence of relationship between the first epidemic burst and rainfall, but a clear relationship between the following epidemic bursts and rainfall after a 3 to 8 day lag.

## 1 Introduction

Wavelet transform has been more and more applied as an effective tool for non-stationary time series analysis when Fourier analysis cannot be used in order to extract the main characteristics of such series [1,2]. These characteristics include periodic variations, regime shifts or sudden perturbations and gaps. Wavelet transform has been used as a signal processing tool in different domains such as climatology [3,4,5], geophysics [6], neurology [7] or image processing [8,9], and for statistical estimation methods [10,11]. More recently, it has been used in epidemiology for analyzing long time series of cases, and as assessing relationships between case time series and factor time series [12-16]. A complete description of wavelet transform analysis can be found in Torrence and Compo [17] and in Nason [18].

Classically, the studied time series are long-term observations (ten years, or more), and present periodic variations that are studied using periodic wavelets. The main information that can be extracted by such analyses are: i) the frequency analysis, assessing periodicity and periodic changes, ii) the time analysis, assessing variations of time and speed, iii) the comparison of two time series (*e.g.* epidemic and climatic series [19]). Wavelet transform analysis is based on the choice of one of the different wavelet functions [20]. This choice is based on the observed characteristics of the time series. Classically, two types of wavelet can be used. The Morlet wavelet function [21] is very useful for frequency analyses of periodic time series [22]. Conversely, the Haar wavelet function is more accurate for time analyses of non-periodic non-stationary time series that exhibit rapid changes [17]. Indeed, frequency and time are related by the uncertainty principle [23]. A wavelet function with accurate frequency localization (such as the Morlet function) can precisely assess frequencies of a periodic time series, but does not present an accurate time localization, leading to less precise analyses of rapid changes. In the case of a recent emergence of a pathogen in a given territory, the lack of historical data precludes from analyzing periodicity (frequency analysis) and leads to analyze time of changes (time analysis).

The Haitian cholera epidemic that began in October 16, 2010 [24] rapidly proved to be the world’s largest epidemic during the Seventh Pandemic [25]. Cholera is an infectious disease due to toxigenic *Vibrio cholera* O1, a bacteria that can be directly transmitted from human to human or via environment. As cholera was imported probably for the first time in Haiti, no historical data could be used to assess periodicity (such as seasonality) even after 2 years. Nevertheless, this short-term time series presented rapid variations that could be explained as different regime shifts or epidemic processes. To accurately extract these times of changes in the epidemic process, a periodic wavelet function (such as Morlet function) was thus considered less useful than a time localized wavelet, such as the Haar function. Yet, the Haar function presents discontinuities that lead to discretization problems [26]. The aim of this work was therefore to develop a continuous wavelet function [27,28], and apply the corresponding wavelet transform to the non-periodic non-stationary time series of the Haitian cholera epidemic.

The present article is organized as follows: Section 2 provides a description of the wavelet transform principles and a brief revue of the different wavelet analysis methods, which highlights the usefulness of the scale and translation parameters. It describes the main properties that should guide the choice of the most suitable wavelet function (continuous and non-periodic). It also presents the new continuous wavelet transform we have developed to assess the non-periodic Haitian cholera epidemic and its relationship with rainfall. Section 3 presents the results of the wavelet transform applied to the cholera epidemic in Haiti. Wavelet spectra of case and rainfall time series were analyzed and cross-wavelet are presented. A comparison to a change point analysis is also presented. Section 4 provides interpretations and discussions of the results.

## 2 Method

### 2.1 Signal analysis

In most cases, the signal analysis process exhibits several phases: the determination of a spectrum, the coding of information, the transmission, and the reconstruction of the signal. For epidemic time series analysis, the whole analysis process is unnecessary: only the determination of spectra at different scales is necessary. Coding, transmission and reconstruction are not useful from an epidemiological point of view. The epidemic records are given as one-dimensional time series and these time series are not stationary. These characteristics are useful to choose an appropriate transform.

The Fourier transform is a decomposition of the signal in order to extract the main frequency components. The Fourier transform requires two conditions for the signal. First, the signal must be stationary [7], *i.e.* with no sudden changes of frequencies and transient periods. Second, it must fit the sum of sinusoids at different frequencies. The Fourier transform does not allow time analysis, and is not applicable in case of sudden frequency variations analysis and scaling change analysis. Moreover, the Fourier integral method considers phenomena in an infinite interval and a spectrum is obtained for the whole signal. This method gives no information on the time localization of a frequency. For all these reasons the Fourier transform cannot be used to analyze a non-periodic non-stationary epidemic time series.

The deficiencies of the Fourier transform in time-frequency analysis were observed by Gabor [23]. In order to study signals having frequencies, which depend on time, the Gabor algorithm provides a localized window function. The signal analysis is performed by a sinusoidal wave of finite duration. The Gabor algorithm defines a time window that divides the sinusoidal wave into finite parts. Between its starting point and its end, the sinusoidal wave has constant amplitude and frequency, and the window is the product of a Gaussian function by a sinusoidal wave. The Gabor function can be written as:

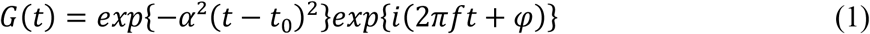

where α, t_0_, f, φ are real constants and *i*^2^ = −1.

The window, centred at *t*_0_, analyzes the corresponding section of the signal series. Then, the window is centred at the following value *t*_1_, closed to the previous one, and so on. An analysis is performed on a first short section of the signal series. Then, the window is shifted and a new analysis is performed. Each section of the signal is analyzed, according to a classical Fourier analysis. The analysis is repeated until the end of the signal series. The frequency of the sinusoidal wave can change when the window is shifted. By introducing a localization parameter *t*_0_ of the section and a frequency *f*_0_, the obtained windowed Fourier transform, also called the Gabor transform, measures locally, around time *t*_0_, the amplitude of the frequency *f*_0_. The signal analysis is a time-frequency analysis. The window width, and consequently the width of a signal section, is arbitrary, and can thus affect the result of the analysis. With such a windowed transform approach, the Gabor transform can be used to analyze non-stationary time-series. Nevertheless, if frequency variations are very fast and with high amplitudes, then the Gabor transform fails to detect them, because the series has to be stationary within each window. The choice of the window width and this piecewise stationary assumption can bias the Gabor transform of time series with sudden changes. Hence, Gabor transform cannot be used for epidemic analysis either.

In order to study signal series showing various scaling properties, Morlet *et al.* [21] have modified the Gabor function to obtain a wavelet. As defined, for the first time, by Grossmann and Morlet [29], the wavelet transform is a decomposition of a signal by translations and dilations of a single fixed function called the mother wavelet Ψ. Wavelet analysis is a time-frequency analysis of the signal, which permits to estimate the spectral characteristics of the signal as a function of time. During the analysis process, wavelet transform can denoise the signal preserving its main features. The sinusoidal functions of the Fourier transform are replaced by pseudo-periodic functions that decay sufficiently fast, oscillating like waves. The mother wavelet integrates to zero, that is:

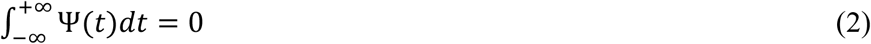

The wavelet transform is well adapted to analyze epidemic time series, providing time-frequency analysis at different scales.

### 2.2 Wavelet properties

Based on a mother wavelet, a wavelet family is obtained by introducing a translation parameter *b* and a scale parameter *a*, and is given by:

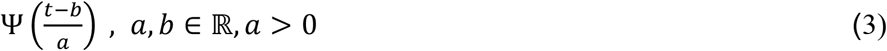

Where ℝ denote the set of real numbers.

The Morlet complex wavelet family can be written as:

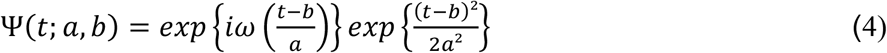

Where *ω* is the angular frequency.

The resolution of a wavelet transform *W* varies with the scale parameter *a*. If the analysing wavelet is narrow, its time localization is precise and accurate, and therefore sudden changes will be detected as well as noise. If the analysing wavelet is wide, the large variations of the signal are exhibited, but the resolution is less accurate and the signal is denoised. The wavelet transform can thus be used as a passband filter, where the bandwidth depends on the value of scale parameter *a* which defined the analysing window width. The smaller the scale parameter *a* (narrow analysing wavelet), the more precise and accurate the time resolution. Defining different values of the scale parameter *a* leads to a multiresolution analysis, which is graphically presented using scalogram figures. Considering the wavelet Ψ as a real function, the wavelet coefficients *W*(*a*, *b*) of a time series *s*(*t*) can be written as:

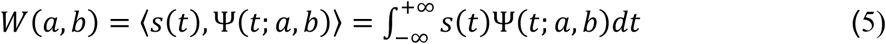

The wavelet coefficients of a time series is defined as the convolution of *s*(*t*) and a translated and scaled version of Ψ(*t*). The time series spectrum is then obtained from *W*, exhibiting the speed of local variations of a time series. To be admissible as a wavelet, a function has to satisfy several crucial properties [29]:

#### Property 1

the function integral converges and is zero mean.

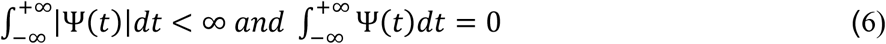

The wavelet values have to divide up, in an equal way, on both sides of the time axis. This implies that Ψ(*t*) oscillates. When this condition is not verified, the positive values (increase of the time series) or the negative values (decrease of the time series) are privileged to the detriment of the other values.

#### Property 2

the function is a square integrable function.

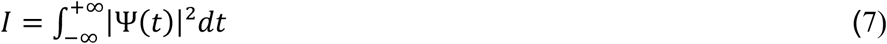

Then, the function is normalized such as

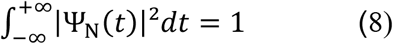

Where Ψ_N_ is the normalized function. This condition is important for being able to compare the results between each other.

#### Property 3

a function is called an orthogonal wavelet if the following condition is verified:

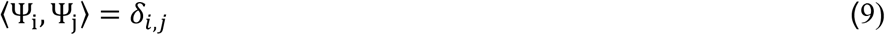

Where *δ*_*i*,*j*_ is the Konecker symbol. This condition is not obligatory, but if it is satisfied the time series decomposition is unique. The wavelet coefficients are not correlated and the time series is a linear combination of the coefficients. A good reconstruction of the original time series can thus be obtained from the wavelet coefficients. The Daubechies wavelets [30,31] form an orthogonal basis and allow a good compression of the information, useful for image analysis. Note that the Morlet wavelet does not satisfy the orthogonality condition.

#### Property 4

moments of the wavelet indicate how a function decays toward infinity. Some vanishing moments could verify the following condition:

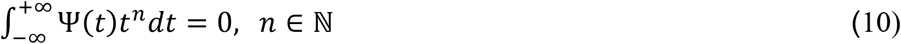

This condition is not obligatory, but leads to calculation simplifications.

Wavelets rarely verify all these properties. It is thus necessary to choose a wavelet which exhibits the properties that are the more useful in the context of the epidemic time series analysis.

### 2.3 Choosing a wavelet for non-periodic series

The choice of a wavelet, for non-stationary time series analysis, is still an open problem. The wavelet function has to reflect the type of features to be extracted. For evident reason, and like many mathematical and statistical approaches, a wavelet must not be chosen only because an algorithm has been developed to perform a wavelet transform. The wavelet function has to be chosen according to the time series to be analyzed, according to the characteristics to be extracted [10,32], and reaching epidemiologic goals (outbreak detection, epidemic mechanism, seasonality, coherence with environmental time series,…).

To choose the right wavelet in our study, we first considered the unnecessary wavelet properties and, second, their mandatory properties to assess epidemic onsets. Reconstruction of the original time series from wavelet coefficients was unnecessary. Therefore, the orthogonality condition (property 3) did not need to be satisfied. In studies of epidemiological time series, the Morlet wavelet has been often used [33-35]. But all these series exhibited periodic cycles. In this context, the Morlet wavelet was a good compromise between the determination of the frequencies of these cycles and the time localization of epidemic bursts. However, the Morlet wavelet should be used with caution because it does not verify property 1, as the mean is not null. In our context, studying frequencies was not of interest since we wanted to localize the epidemic bursts. The uncertainty relation implies that a good accuracy of the frequency values leads to a less accurate identification of time localization and conversely. As good accuracy of the time localization was needed in our context, the Morlet wavelet was not chosen. For the same reason we did not choose the Shannon wavelet [18].

Mandatory properties of wavelets for our study were related to the following. We studied a new cholera epidemic over a relatively short period (about 800 days). Epidemiological series was not smooth and exhibited many sudden transitions of large magnitude. The lack of annual period made it unnecessary to have a wavelet defined on the complex set ℂ, but to use only a real wavelet. Our goal was to precisely date the epidemic bursts. Thus, a wavelet well localized in time was needed to obtain a good resolution. To choose the mother wavelet we kept the following properties:

– Properties 1: the wavelet integral must be convergent and zero mean,
– Property 2: the wavelet must be normalized.

Moreover, the wavelet had to decay sufficiently fast. The wavelet that best satisfies these properties is the Haar wavelet [18,36]. It is real and precisely localized in time. Its main drawback is that this wavelet is a piecewise-defined function and it is not regular. It presents several significant discontinuities with substantial angularities, which may interfere with the practical computation, especially with a discretization in the vicinity of a discontinuity [26]. Hence, this wavelet may be suitable for the analysis of regular functions but not for sudden changes in time such as epidemic time series. Therefore, we chose a smooth and continuous wavelet, with a shape similar to the Haar wavelet. This chosen mother wavelet, called GCP wavelet, was established by Gaudart et al. [28], and is defined as:

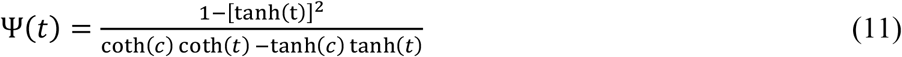

where tanh and coth are hyperbolic functions.

The value of the parameter *c* ∈ ℝ fixes the slope of the wavelet flanks. Depending on the value of this parameter the increase and decrease of the curve is more or less rapid, setting the decay of the wavelet. In the following, *c* will be fixed to 1, unless special statement. This value allowed a rapid return to zero, a sufficiently fast decrease of the wavelet, and a precise time localization. Too large values of *c* led to a slow decay and therefore a poor extraction of sudden changes in the epidemic time series. Small values led to discontinuities.

In what follows, the function Ψ(*t*) represents the GCP wavelet, unless otherwise indicated. This function Ψ is defined on ℝ, convergent, zero mean, and it is a square integrable function. Consequently, the function Ψ is a wavelet. The GCP wavelet family Ψ(t; a, b) is given by:

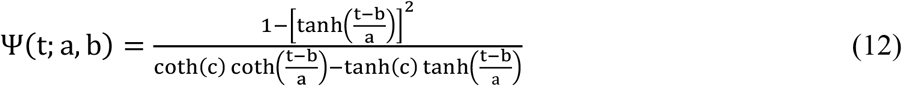

with *a* ∈ ℝ^+*^, *b* ∈ ℝ, *c* ∈ ℝ*.

In order to compare wavelet transforms between each other, the wavelet Ψ(*t*; *a*, *b*) needs to be normalized [28], following equation (8). The normalized wavelet Ψ_N_(t) is written as:

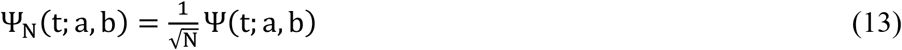

Where *N* is the normalized factor such as:

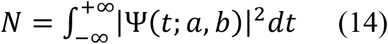

The norm depends on the scale parameter *a*. In the development of the algorithm, the norm *N* thus has to be calculated for each value of the scale parameter. Unless special statement, Ψ(*t*) will be assumed to be the normalized GCP wavelet.

In our context, the analysis of the epidemic time series was performed using wavelet translation, whatever the width of the wavelet. The entire epidemiologic time series was scanned by the wavelet, including a period prior the beginning of the cholera epidemic. Therefore, the value of the translation parameter *b* ranged from 1 to 812 days. Variation of the scale parameter showed different wavelet shapes (figure 1). The upper curve in figure 1 is a narrow wavelet (scale parameter *a* = 1), centred at *t* = 5 (translation parameter *b*= 5). The middle curve is a wavelet of average width (*a* = 2), centred at *t* = 7. The lower curve is a wide wavelet (*a* = 3), centred at *t* = 9. In order to determine the wavelet width, we defined, for a given time *t*, the ratio:

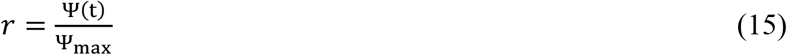

Where Ψ_max_ > 0 is the maximal positive value of the wavelet for a given value of the scale parameter. Width of the useful part of a wavelet is given by the condition:

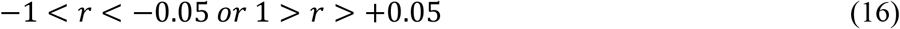

**Figure 1.**
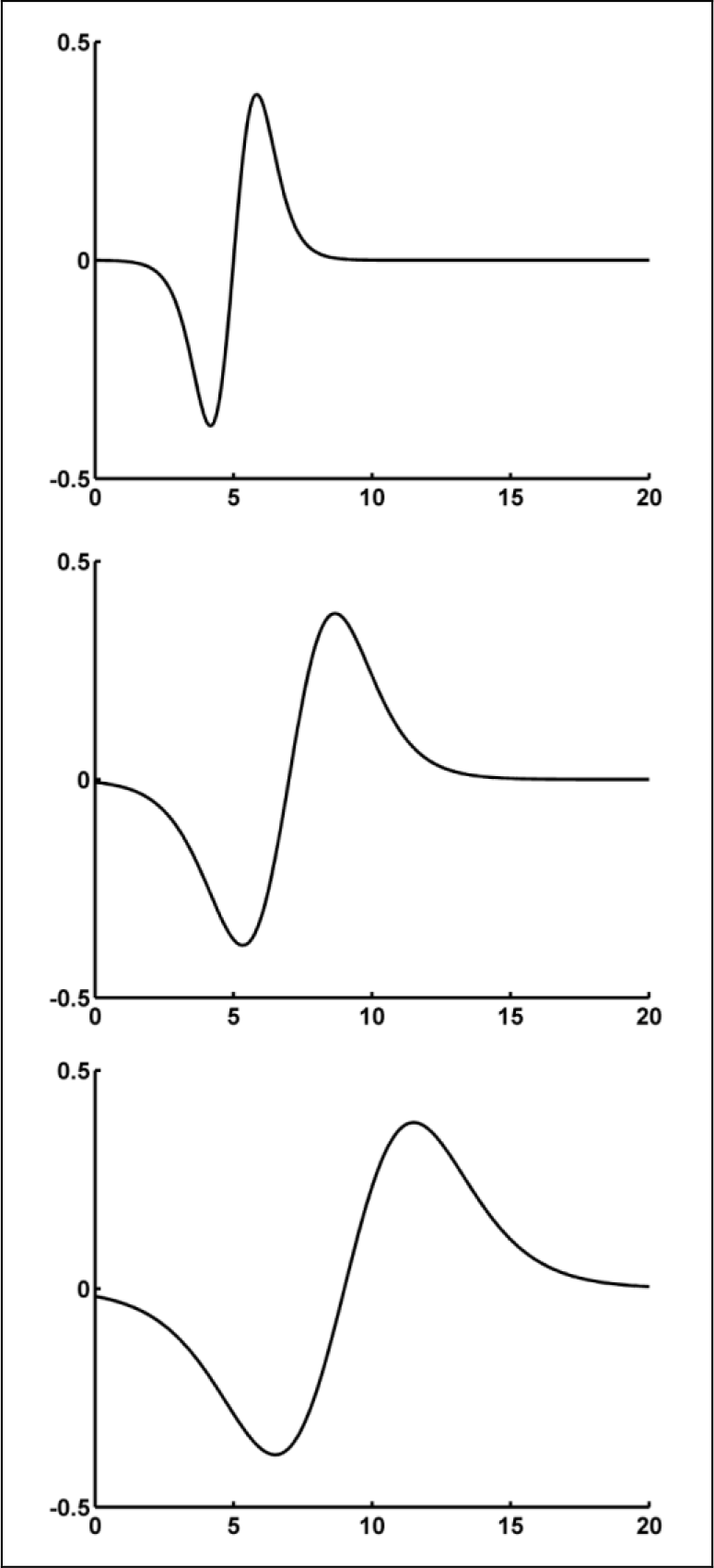
Wavelet variations as a function of time. Non normalized wavelets Ψ(*t*; *a*, *b*) are plotted for different values of scale and translation parameters. The x-axis represents the time (days). The y-unit is arbitrary. Upper panel: *a* = 1, *b* = 5; central panel: *a* = 2, *b* = 7; lower panel: *a* = 3, *b* = 9.

Outside these ranges the values of Ψ(*t*) are negligible. Using this relation, the scale parameter can be interpreted as the wavelet width in time units (here in days) (table 1).

**Table 1.**
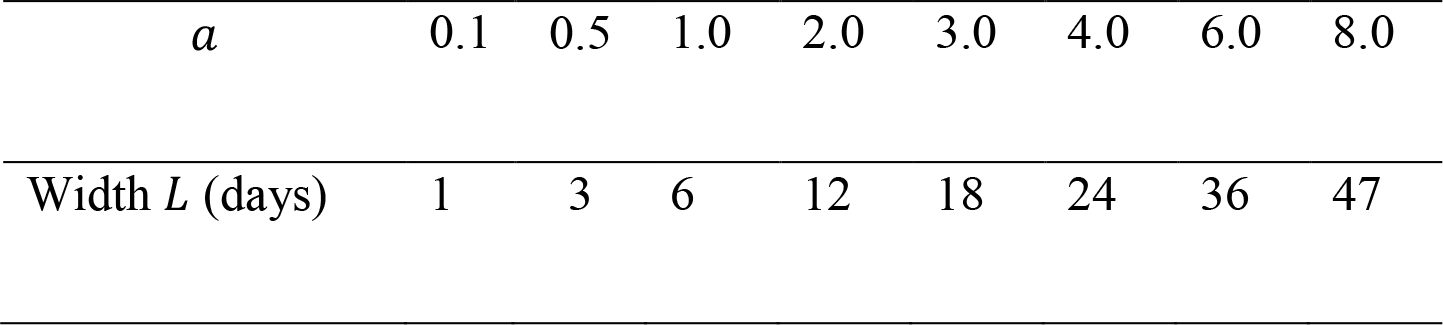
Relationship between the scale parameter *a* and the observed time width *L* of the wavelet (days).

For a value of *a* = 0.1, all daily changes were analyzed in detail by the wavelet. For a value of *a* = 6, the time series was smoothed and we obtained the main tendency of changes through about 36 days. Higher values of the scale parameter *a* smoothed all time variations of the series, with no interest from an epidemiological point of view. Hence, for a study of about 800 days, the useful range of the parameter *a* was between 0.1 and 6.0. In the context of longer time series and/or slower disease transmission, larger values of the scale parameter may be more suitable.

### 2.4 Wavelet spectrum and cross-wavelet spectrum

Real wavelet transform can highlight the local characteristics of a time series, in particular its rapid changes and burst amplitudes. Let us consider the epidemiological series *s*(*t*) and the wavelet coefficients *W*(*a*, *b*). The local power spectrum is defined as:

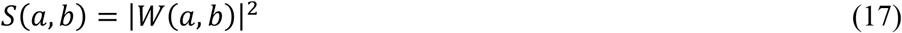

For a given scale, that is a given resolution, the spectrum varies as a function of the translation parameter. These variations exhibit the spectral peaks of the original time series. The magnitude of each peak represents the epidemic series dynamic: the more rapid the variation, the higher the spectral peak. A nonzero constant time series has a null spectrum. In epidemic analysis, it is of high interest to note the increase or decrease of case toll, but, according to equation (17), spectra have only positive values. For this interpretability reason, we modified the definition of the spectra as follows:

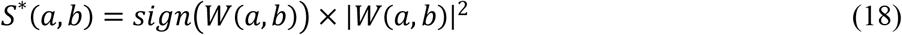

Thus, an increase of the time series at time *t* was characterized by a positive value of *S*^*^(*a*, *b*) at *b* = *t* (the faster the increase, the higher value the spectrum). Conversely, a decrease of the time series at *t* was characterized by a negative value of *S*^*^(*a*, *b*) at *b* = *t* (the faster the decrease, the lower value the spectrum). Absence of change in time series provided a null spectrum, whatever the cholera case toll.

As rainfall is an important driver of cholera epidemic, we also analyzed rainfall time series, and compared epidemic wavelet spectra and rainfall wavelet spectra. The wavelet coherence, measuring the cross-correlation between two time series, has been defined for complex wavelets [17,22]. In the case of real wavelets, such as the GCP wavelet, the wavelet coherence is equal to 1 whatever times and scales, and therefore cannot be used. Therefore, we used the cross-wavelet spectrum defined by the following equation:

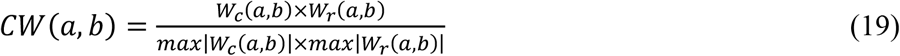

where *W*_*c*_(*a*, *b*) and *W*_*r*_(*a*, *b*) are the cholera epidemic coefficients and rainfall coefficients, respectively. A denominator was added to the classical cross-wavelet spectrum definition in order to re-scale both spectra at a same magnitude.

Cross-wavelet spectra reveal the underlying relationship between two time series. In our study it highlighted the interactions between rainfall and cholera epidemic. Furthermore, to provide a thorough description of these relationships, a lag parameter between rainfall and cholera cases was introduced in the cross-wavelet spectrum. By analyzing cholera cases at a time *t* and rainfall at a time *t*-lag, the lag parameter varied from 0 to a value specified when a cross-wavelet maximum was reached. These lags indicated the number of days between rainfall and its potential consequence on the cholera case toll. In order to evaluate cross-wavelet at different lags and times, lagograms were developed that provided cross-wavelet spectra varying through lags and times.

Program development and data analysis were conducted using Matlab 7.0.1 (The Mathworks Inc., Natick, MA, USA).

### 2.5 Comparison to a change point analysis

Results obtained by the developed wavelet analysis of the cholera case toll have been compared to a change point analysis, applying the Segment Neighbourhood (SN) method [37]. This method is considered as an exact one for change point detection, evaluating all possible segmentations to a maximum number of change points. This maximal number of change points, 14, have been chosen higher than the number of spectral peaks determined by the wavelet analysis. A Poisson distribution and a penalty value of 1.5 × *log*(*n*) have been applied. Change detection in mean and variance was performed using R^®^ v3.0.1 (The R Foundation for Statistical Computing, Vienna, Austria) with the *changepoint* package, provided by Killick et al. [38].

## 3 Application

### 3.1 Cholera time series analysis

During the study period, the daily cholera case toll was prospectively collected by Haitian Departmental Health Directorates. Departmental databases were sent to the Haitian Directorate of Epidemiology, Laboratory and Research (DELR), where data were gathered and analyzed after quality control [24,25]. In this study, we did not include personal data but only the national daily toll of cholera cases. Its time series was analyzed from September 01, 2010 to November 20, 2012 (812 days). In order to eliminate edge effects, we excluded from the results the 10 first days and the 10 last days. Calculation of the wavelet coefficients (using the normalized wavelet and scanning the whole period) provided a spectral value for each value of the translation parameter *b*, from *t* = 1 to 812 days, and for each value of the scale parameter *a*.

The time series and spectra (at different values of the scale parameter *a*) are shown in figure 2. The scalogram represents spectrum values according to the scale and the translation parameters (figure 2B).

**Figure 2.**
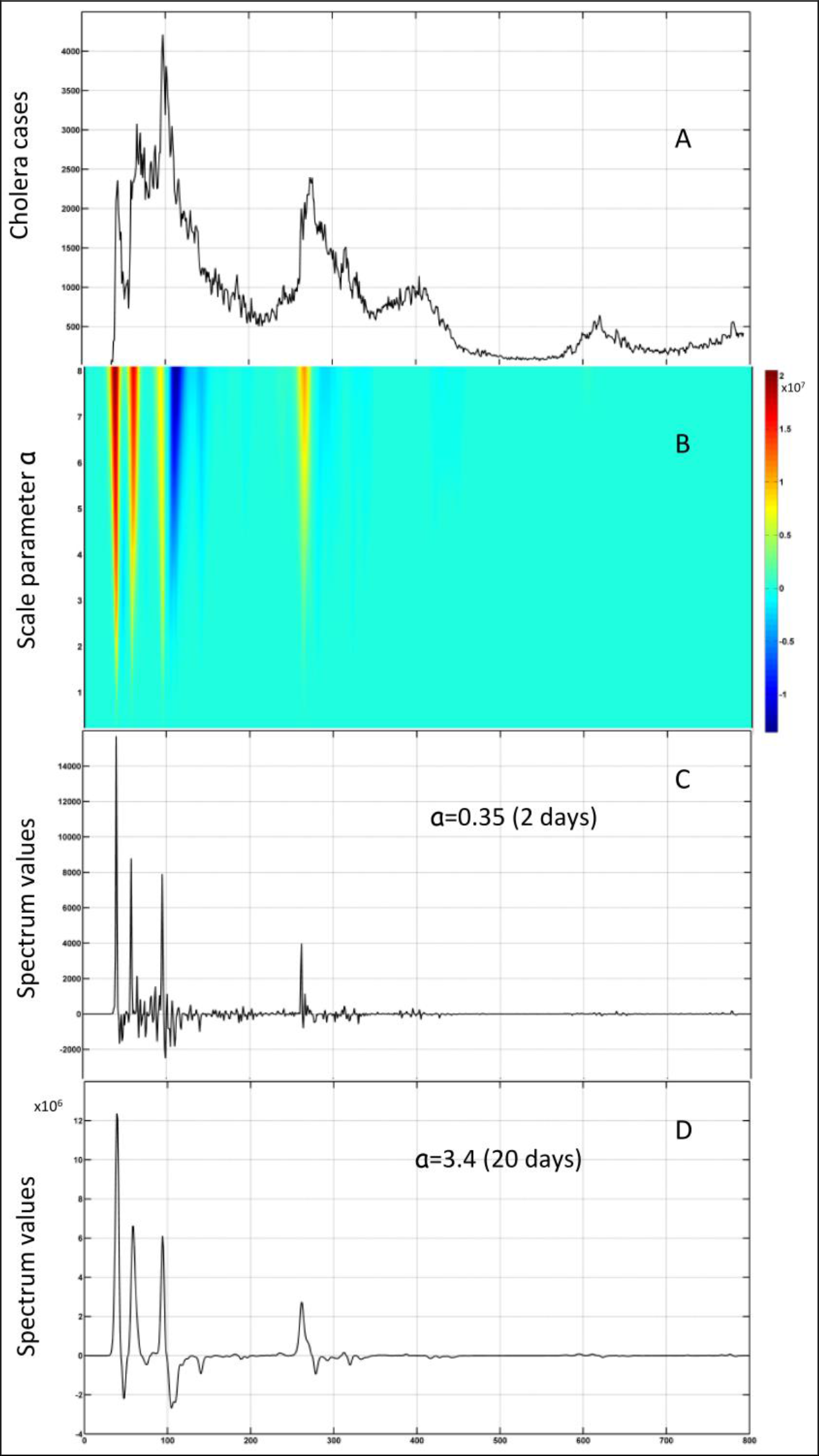
Cholera time series and wavelet transform. The x-axes represent the time (days) from September 10, 2010 to November 10, 2012. A) Daily reported cholera case toll; B) Cholera case toll scalogram (*a* from 0.1 to 8). The coloured scale represents the values of the spectra *S*\*(*a*, *b*) (arbitrary units). High values are shown in dark red and low values (negative) in dark blue; C) Spectra for a narrow wavelet width: *a*=0.35; D) Spectra for a large wavelet width: *a*=3.4.

The spectral analysis (figures 2C and 2D) highlighted 4 spectrum peaks corresponding to 4 bursts of the epidemic that were related to different epidemic mechanisms. The first spectrum peak corresponded to the explosive start at the lower Artibonite valley on October 19, 2010 (just after its initiation in the commune of Mirebalais [24]). This high spectrum peak had an impact towards the whole study period, as it remained high at large values of the scale parameter. The second spectrum peak, with a lower magnitude, corresponded to the epidemic spread out of the Artibonite valley (around November 07, 2010). The third spectrum peak corresponded to worsening in cholera incidence concomitant to riots in Port-au-Prince following the first round of presidential elections in early December 2010. This spectrum peak was followed by negative spectrum values mainly shown at large scale (figure 2D), corresponding to the slow decrease of the epidemic. After a lull phase, the fourth spectrum peak corresponded to the rainy season (May 2011). This peak was related to an outbreak that showed a profile different from the first ones: its dynamic was more progressive [25]. It was highlighted by the lower magnitude but larger area under the curve of the fourth spectrum peak when considering large values of the scale parameter. The magnitude of these 4 spectrum peaks still remained even if using a larger scale parameter (*a* = 8). Following these four peaks, the magnitude of the spectrum decreased to undetectable spectrum values which can be interpreted as a slow trend towards epidemic stability as time passed. The particularly high magnitude of the first spectrum peak explained the major part of the variability of the time series. Subsequent small variations of the cholera case toll could thus not be seen.

The change point analysis (figure 3) detected a high number of change points (q=14): the epidemic onset and the 2 following peaks have been detected, as well as the increases in June 2011 and May 2012. But the decrease at the end of October 2010 has not been detected. The decrease between December 2010 and March 2011 has been split in 3 segments, as well as the decrease from June to August 2011.

**Figure 3.**
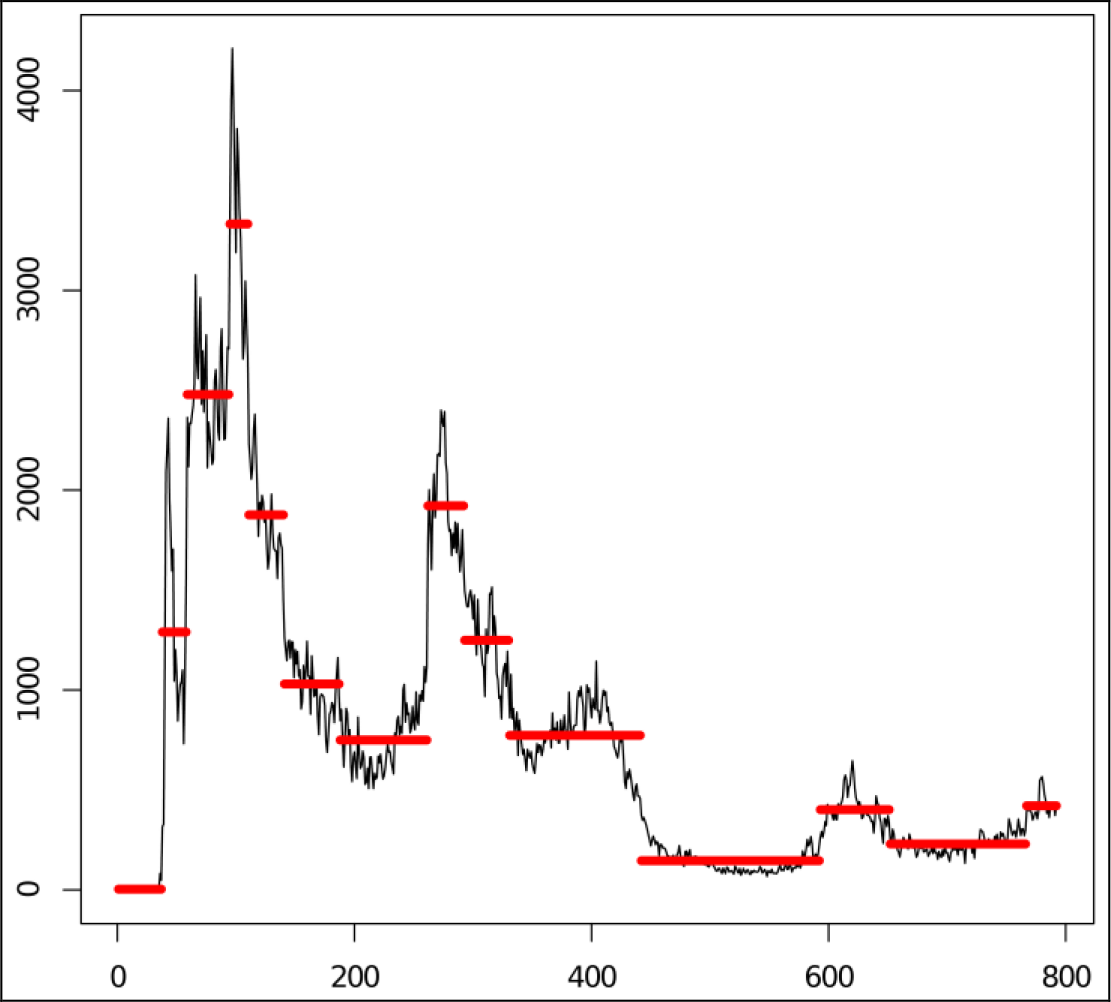
Cholera time series and Change Point Analysis. The x-axis represents the time (days) from September 10, 2010 to November 10, 2012. The y-axis represents the Cholera case toll. The red horizontal lines identify the changes in the cholera case mean.

### 3.2 Rainfall time series analysis

Daily accumulated rainfall data were obtained through the NASA Goddard Earth Sciences, at the national level. These observations were derived from the Tropical Rainfall Measuring Mission (TMPA-RT 3B42RT); for more details see http://disc.sci.gsfc.nasa.gov/giovanni/overview/index.html (access: September, 2013). Rainfall time series and spectra (at different values of the scale parameter) are presented in figure 4. The scalogram represents spectrum values according to the scale and translation parameters (figure 4B). The spectral analysis of rainfall exhibited two major spectrum peaks related to hurricane Tomas (November 05, 2010) and hurricane Sandy (October 24, 2012), respectively. Peaks detected in the meanwhile, especially for large values of the scale parameter, corresponded to the 2011 and 2012 rainy seasons. No high rainfall had been exhibited before the first cholera case burst.

**Figure 4.**
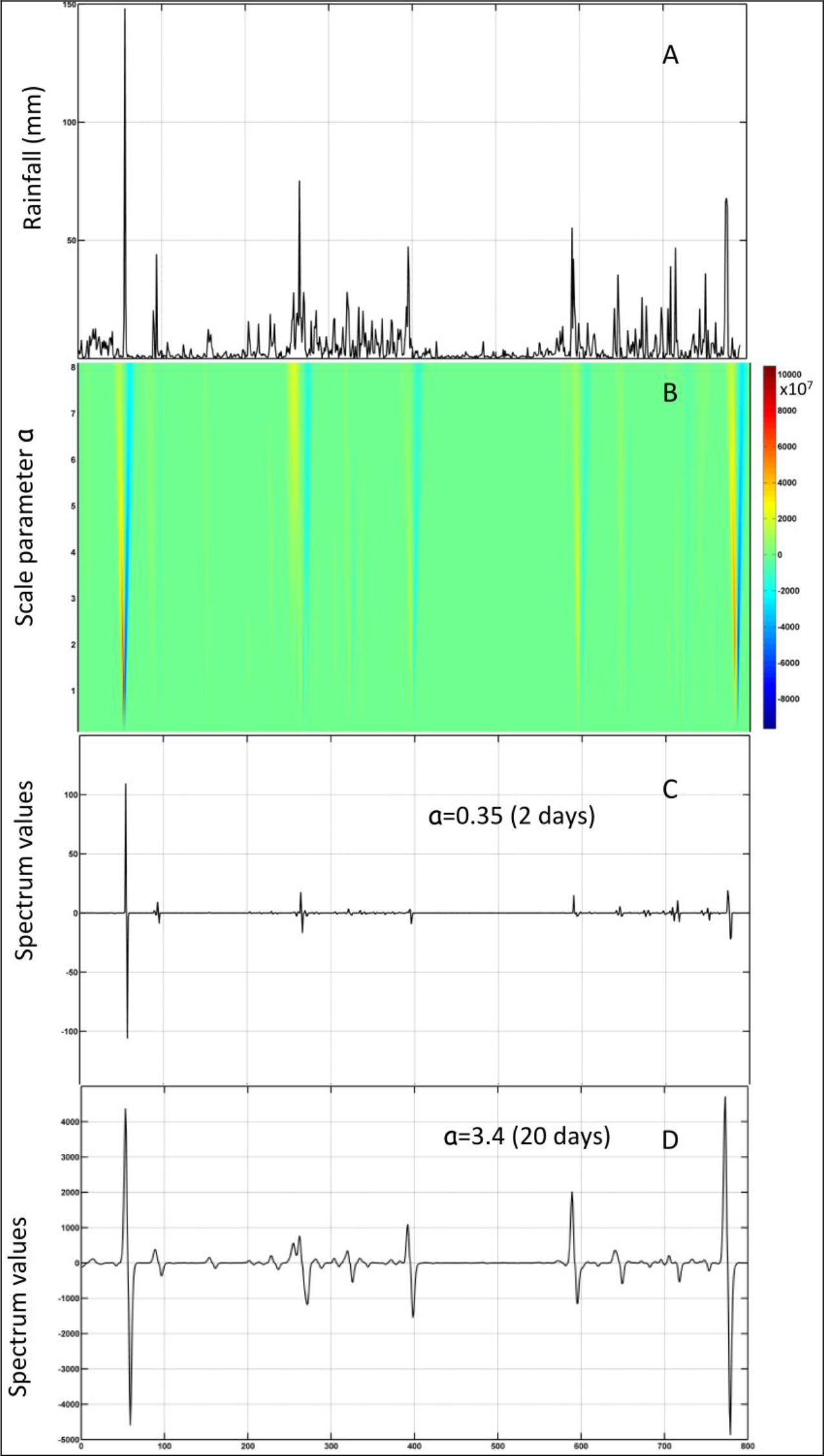
Rainfall time series and wavelet transform. The x-axes represent the time (days) from September 10, 2010 to November 10, 2012. A) Daily reported rainfall; B) Rainfall scalogram (*a* from 0.1 to 8). The coloured scale represents the values of the spectra *S*\*(*a*, *b*) (arbitrary units). High values are shown in dark red and low values (negative) in dark blue; C) Spectra for a narrow wavelet width: *a* = 0.35; D) Spectra for a large wavelet width: *a* = 3.4.

### 3.3 Cross-wavelet spectrum

The cross-wavelet spectra calculated with a null time lag are presented in figure 5. They showed a globally low relationship between cholera cases and rainfall, except for the epidemic on May 2011 at large values of the scale parameter (figure 5C), what field epidemiologists also reported. The negative values of the cross-wavelet spectra at the beginning of the studied period can be interpreted as a decoupling of the 2 time series. The relationship between the two time series had thus to be studied at different time lags. Indeed, a lag around 3-4 days, between rainfall and cholera case toll is highlighted by the lagogram, for scale parameter *a* = 1.7 (10 days) (figure 6). Maximum of the cross-wavelet spectra occurred for a 3 day lag at *b* = 58 days, around November 08, 2010 (second cholera case burst), 3 days after hurricane Tomas reached Haiti (table 2). Concerning the third and fourth cholera burst, a small amount of rainfall was reported 5-6 days before the third cholera burst (around December 15, 2010). The rainy season began around May 20, 2011, 5-6 days before the onset of the fourth cholera case burst. It is noticeable that no relationship had been exhibited between the decrease of both the cholera case toll and rainfall. Furthermore, high cross-spectra were followed by very low (negative) cross-spectra (figure 6), i.e. the transient coupling of the two series were systematically followed by their decoupling. This can be interpreted as a booster effect of rainfall on an ongoing cholera epidemic, after which transmission is neither further accelerated by persisting rainfall, nor immediately stopped with rain cessation.

**Figure 5.**
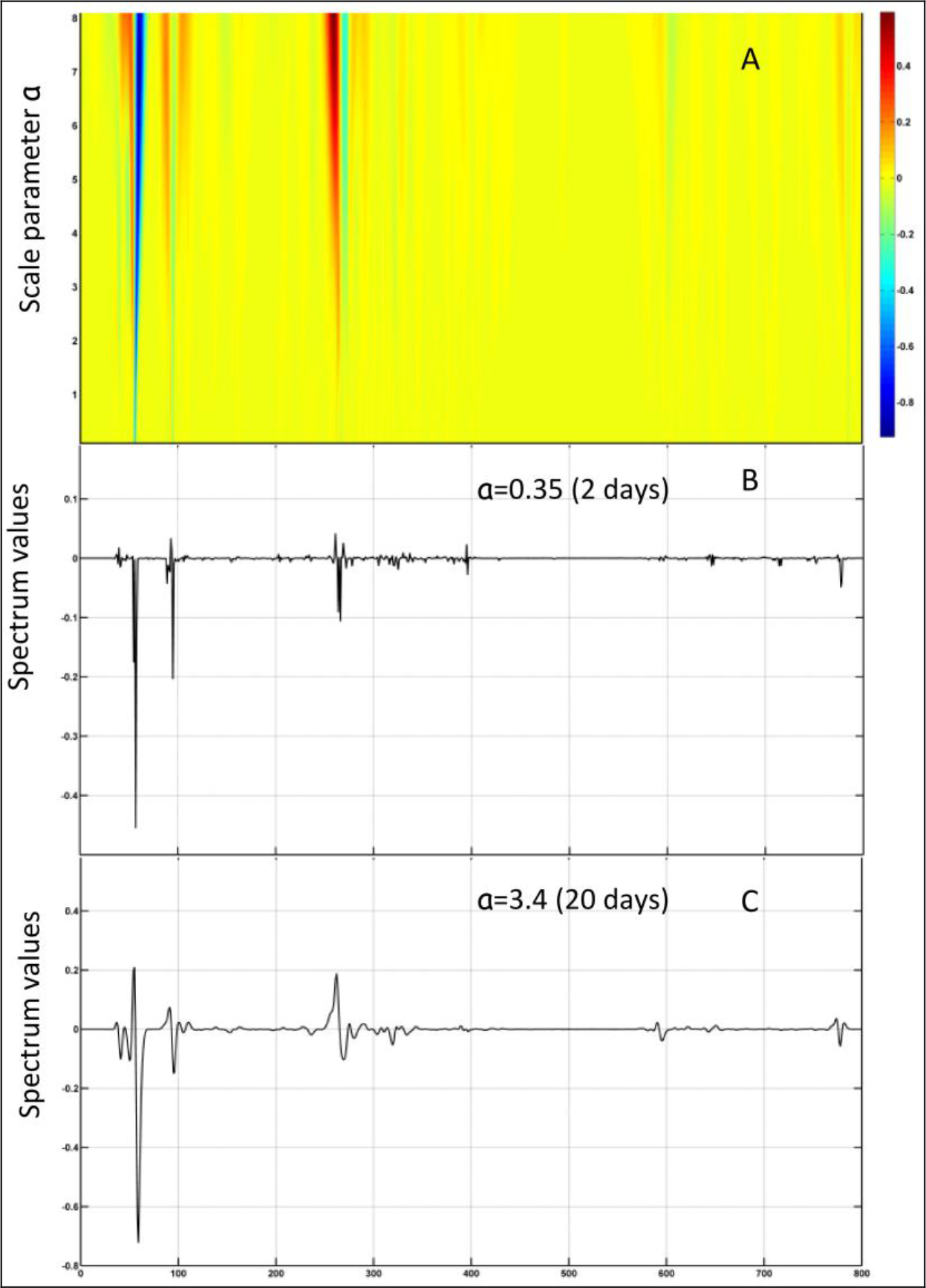
Cholera case toll and rainfall cross-wavelet spectra, lag=0. The x-axes represent the time (days) from September 10, 2010 to November 10, 2012. A) Cross-wavelet scalogram (*a* from 0.1 to 8). The coloured scale represents the values of the cross-spectra *CW*(*a*, *b*) (arbitrary units). High values are shown in dark red and low values (negative) in dark blue; B) Cross-spectra for a narrow wavelet width: *a* = 0.35; C) Cross-spectra for a large wavelet width: *a* = 3.4.

**Figure 6.**
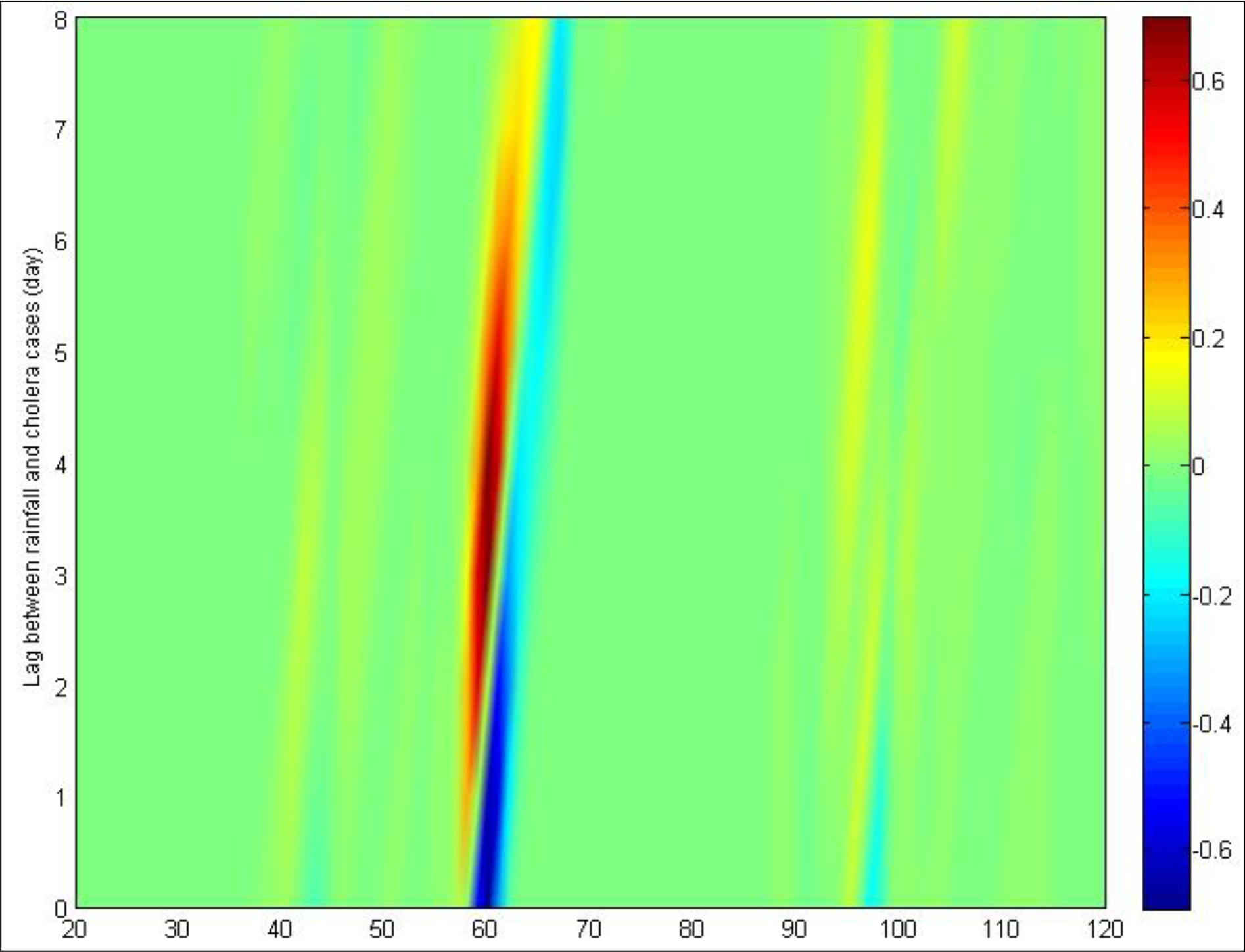
Cross-wavelet lagogram: cholera case toll and rainfall cross-wavelet spectra, with various lags; *a* = 1.7 (10 days). The x-axis represents the time (days) from October 01, 2010 to January 08, 2011 (100 days). The y-axis represents the value of the lag between rainfall and cholera case toll, ranging from 0 to 8 days. The coloured scale represents the values of the cross-spectra *CW*(*a*, *b*) (arbitrary units). High values are shown in dark red and low values (negative) in dark blue.

**Table 2.** Cross-wavelet spectrum maxima. Maximal values of cross-spectra are presented according to the lag between cholera case toll and rainfall, for *a* = 1.7 (10 days wavelet width). The time column represents the related date of the cholera case toll.

| Lag (days) | Cross-spectrum maxima (a.u.*) | Time<br>(days from Sept. 10, 2010) |
| --- | --- | --- |
| 1 | 0.2684 | 56 |
| 2 | 0.5601 | 57 |
| 3 | 0.6956 | 58** |
| 4 | 0.6766 | 58 |
| 5 | 0.5513 | 59 |
| 6 | 0.3377 | 60 |
| 7 | 0.2042 | 62 |
| 8 | 0.1729 | 63 |

\* a.u.: arbitrary unit
\*\*November 08, 2010

## 4 Discussion

Applied to non-periodic non-stationary epidemic time series, the wavelet transform, using the GCP wavelet, highlighted the main features of the cholera epidemic in Haiti and its relationship with rainfall. The wavelet spectra provided timely accurate information about speed and intensity of epidemic variations. These variations could not be easily evaluated by only observing the case toll time series, because of the violence of the first bursts due to the particular context of this epidemic. Wavelet transform also highlighted slower events such as the epidemic decreases after the third and the fourth bursts, in a multi-resolution approach. As spectrum calculation was based on a convolution between the time series and the wavelet, each spectrum was calculated considering the whole time series, i.e. the whole epidemic dynamic. Using a continuous scale parameter, the wavelet transform exhibited the epidemic variations at different time scale, as showed by scalograms. This was of high interest for field epidemiologists, as it led to more accurately assess the epidemic evolutions. Three types of spectral peaks have been observed: i) peaks only observed at narrow scale values, which were very precisely localized in time; ii) peaks only observed at large scale values, which exhibited slower epidemic dynamics through larger time intervals; iii) peaks observed both at narrow and large scale values, which not only exhibited a time localized variation, but also a variation that influenced a large time interval. Persistence of high wavelet spectra along a large range of scale parameter values highlighted the magnitude of the first burst, and its impact on the whole epidemic.

The formal relationship between wavelet approach and statistical analysis has been made only for stationary time series under the assumption of dyadic sample size [18,39]. The spectrum expectation can be interpreted as the variance of the time series although this has not been yet formally demonstrated for non-stationary time series and non-dyadic sample size. Nevertheless, for a given scale parameter *a*, the spectrum expectation can be interpreted as a variability measure of the time series. This has practical implication in interpreting the spectra. Indeed, the relative importance of a variation at time *t*, relatively to the whole time series, can be measured by the relative magnitude of the spectrum value for *b* = *t*, relatively to the spectrum expectation. From an epidemiological point of view, the case spectra need a separate evaluation for the second year of the epidemic. Indeed, as relative measures, spectra of the second year are much smaller than the 4 peaks of the first year. The second year counted for about 102,582 cases (vs 493,069 cases during the first year), and cholera outbreaks of lower magnitude appeared during the rainy seasons (around October 2011 and May 2012).

In our context, the choice of the GCP wavelet has been mainly led by the non-stationary feature and the non-periodicity of the case and rainfall time series, and by the research of precise time localization of their features. This wavelet with a null mean and no period presents one positive and one negative part only, both with equal areas under the curve. With its simple analytical form, the GCP wavelet led us to quantify the rapid changes of the cholera case toll. The parameter *c* (see equation 11), which remained fixed in this study, can be used to tune the speed of the steady-state wavelet return.

Spectral analysis, through wavelet transform, can be considered as an alternative to the classic change point analysis [37,38]. Wavelet approaches provide different information about both time and frequency domains. It allows to extract the main features explaining the behavior underlying a time series, facilitate aberration detection and epidemiologic analysis. In our context, a few features (timely localized or periodic) explaining a time series can be useful from an epidemiological point of view, as well as studying relationships between two time series. Classical change point analyses are considered as linear approaches and do not allow relationship analysis. As a nonparametric non-linear approach, wavelet analysis is able to adapt to local and global features, which is not possible with classical linear approaches [10]. GCP wavelet approach could be particularly interesting to analyze disease surveillance time series.

## 5 Acknowledgements

The authors acknowledge Anestis Antoniadis for his review of this manuscript and Louis Gaudart for his helpful comments on wavelet theory, particularly on the GCP wavelet. The authors also acknowledge Dr Roc Magloire, from the MSPP, Haiti.

### Funding statement

The epidemiological field study was supported by the French Embassy in Port-au-Prince (Haiti). Dr. Jean Gaudart was also supported by the ADEREM association for biological and medical research development (Association pour le Développement des Recherches biologiques et Médicales, http://www.aderem.fr). The funders played no role in study design, data collection and analysis, the decision to publish, or preparation of the manuscript.

### Competing interests

The authors declare that no competing interests exist.

## 6 Data accessibility

Cholera case time series is available through the website of the Ministry of Health of Haiti (http://mspp.gouv.ht/newsite/). Rainfall time series is available through the NASA Goddard Earth Sciences website (http://disc.sci.gsfc.nasa.gov/giovanni/overview/index.html).

